# “Midi-metagenomics”: A novel approach for cultivation independent microbial genome reconstruction from environmental samples

**DOI:** 10.1101/2023.01.26.525644

**Authors:** John Vollmers, Maximiano Correa Cassal, Anne-Kristin Kaster

## Abstract

Since the majority of microbial organisms still evade cultivation attempts, genomic insights into many taxa are limited to cultivation-independent approaches. However, current methods of metagenomics and single cell genome sequencing have individual drawbacks, which can limit the quality as well as completeness of the reconstructed genomes. Current attempts to combine both approaches still use amplification techniques which are prone to bias. Here, we propose a novel approach for the purpose of genome reconstructions, that utilizes the potential of cell sorting for targeted enrichment and depletion of different cell types to create distinct cell fractions of sufficient size, circumventing amplification. By distributing sequencing efforts over these fractions as well as the original sample, co-assemblies become highly optimized for co-abundance variation based binning approaches. “Midi-metagenomics” enables accurate metagenome assembled genome (MAG) reconstruction from individual sorted samples with higher quality than co-assembly of multiple distinct samples and has potential for the targeted enrichment and sequencing of microbial dark matter.

## Introduction

According to current estimates, less than 1% of environmental prokaryotes are culturable under laboratory conditions^1,2^. The vast majority of microorganisms currently evade direct analysis with classic microbiological methods and is thus commonly referred to as ‘microbial dark matter’^1,3^ (MDM). However, advances in cultivation-independent methodologies such as metagenomics and single-cell genomics nowadays enable characterization of uncultured organisms^4–6^ **(Figure 1)**.

**Figure 1:**
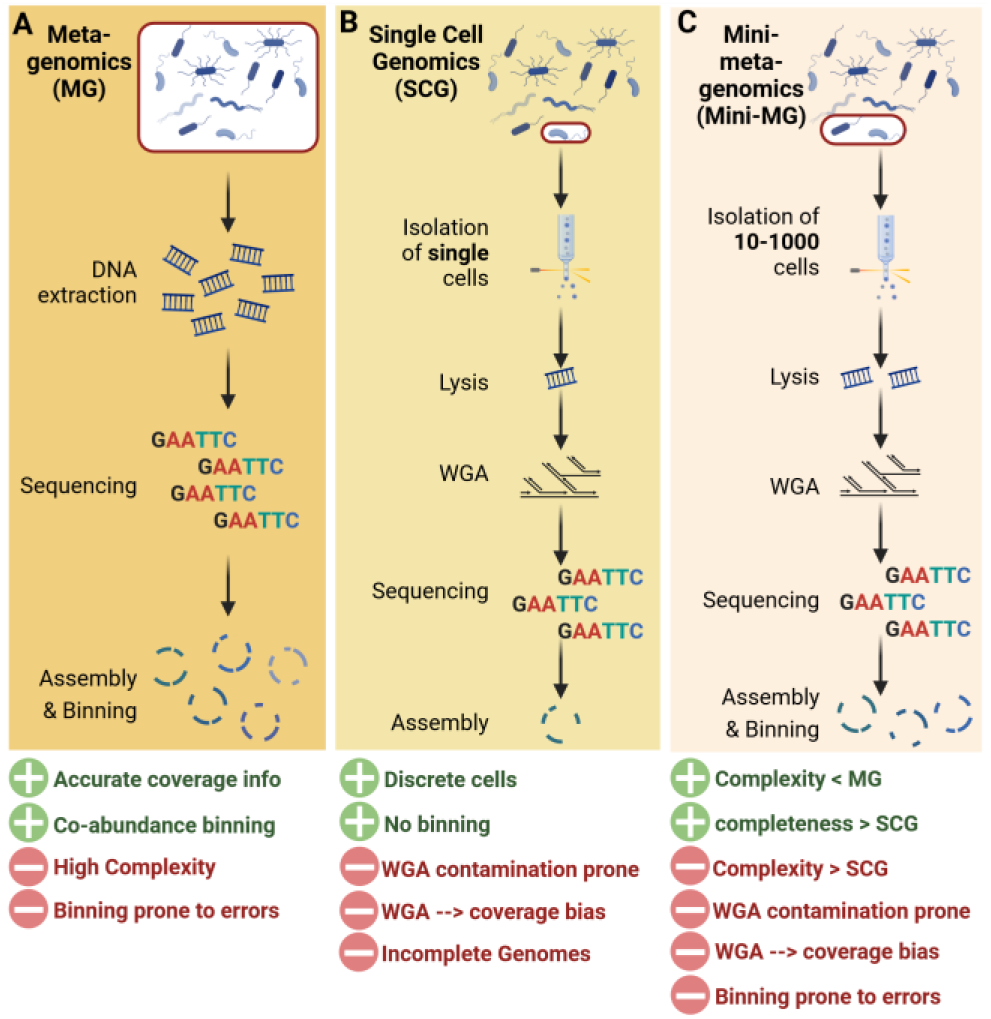
Current culture-independent methodologies. In (**A**) Meta-genomics the entire DNA of an environmental community is sequenced. Assembled contigs are binned into metagenome assembled genomes (MAGs). In (**B**) single-cell genomics (SCG), individual cells are isolated, sequenced and analyzed. Due to little DNA content per cell, whole genome amplification (WGA) is required. In (C) Mini-metagenomics typically pools of 5-1000 cells are sortied and sequenced. While complexity is lower than for standard metagenomics, binning is still required, and the low DNA content of small cell pools still necessitates WGA. Created with BioRender.com

In metagenomics **(Figure 1A)**, the entire DNA of an environmental community is extracted, sequenced, and analysed^7^. Unfortunately, the reconstruction of individual discrete genomes is sometimes not possible, especially for highly complex communities and organisms of low abundance. As a result, metagenome assembled genomes (MAGs) are highly susceptible to chimerism, meaning that they can contain contigs that originate from the genomes of different taxa^8,9^.

Single-cell genomics (SCG) **(Figure 1B)** circumvents this risk by targeting individual cells^10,11^. However, since a single prokaryotic cell contains only a few femtograms of DNA and the minimum requirement for high throughput sequencing is typically in the nanogram range, a whole genome amplification (WGA) is required^12,13^. This is a severe disadvantage, as WGA usually yields extremely uneven read coverage, constituting bias that is particularly pronounced for genomes with high GC content and usually results in fragmented as well as incomplete single cell amplified genomes (SAGs)^11,14,15^.

In order to minimize these drawbacks and maximize the advantages of both methods, there is a strong interest in combining single-cell and metagenomic approaches. A current example for such an attempt is “mini-metagenomics”^16^, which targets small groups of usually 5-100 cells **(Figure 1C)**. These cells are then sequenced together and subsequently treated as a simplified metagenome^14,16^. The DNA yield of such small cell groups is, however, still not sufficient to circumvent amplification, but is thought to efficiently reduce random bias. Furthermore, the relatively low complexity of such mini-metagenomes should, in theory, allow for better genome reconstructions than the more complex metagenome of the original community. However, this approach is still affected by systematic WGA bias that may be caused by e.g. variations in GC content^17^. Most importantly though, effective binning criteria are limited because contig abundance information is not available due to the uneven read coverage, a severe drawback that also obstructs the currently most effective binning strategy: co-abundance variation across samples^18^. Therefore contigs will likely have to be binned exclusively based on nucleotide signatures, which is less reliable, especially for short contigs of highly fragmented genomes^9^.

We here present an alternative approach, termed ‘midi-metagenomics’, that utilizes cell sorting to create custom community fractions of sufficient cell count to circumvent the need for amplification entirely. Fluorscence-activated cell sorting (FACS) is used for targeted enrichment and depletion of different cell types to create fractions which are highly optimized for co-abundance variation based binning approaches. This way, the quality of genome reconstructions can be maximized, even if only individual samples without spatial or temporal parallels are available **(Figure 2)**.

**Figure 2:**
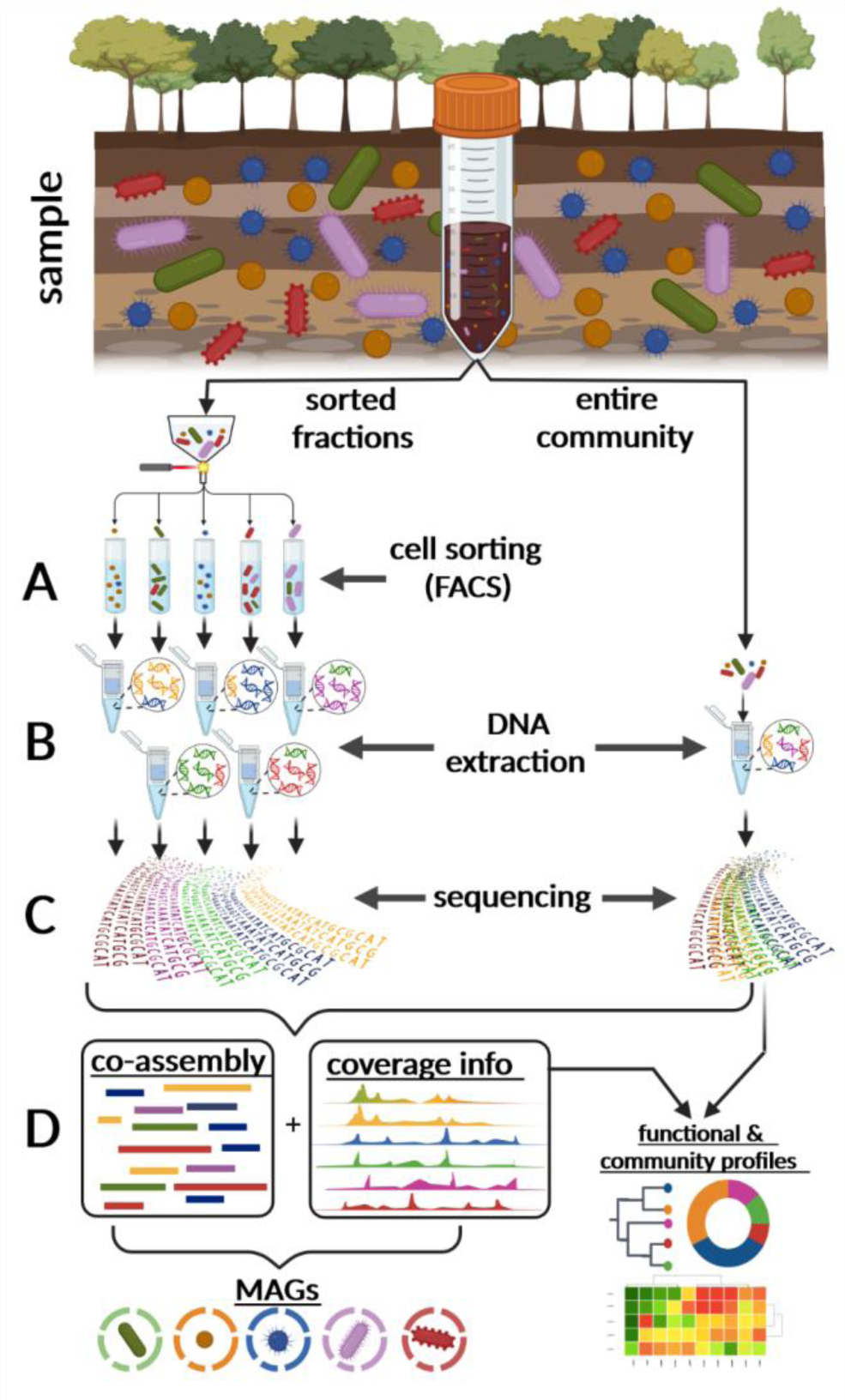
Midi-metagenomics workflow. **(A)** Part of the sample community is fractionated into distinct groups of several hundred thousand to millions of cells by cell sorting. Different cell types are not separated with absolute stringency, but differentially enriched **(B)** DNA is extracted separately from each fraction, as well as the original unsorted sample. **(C)** Extracted DNA is sequenced directly without whole genome amplification (WGA). **(D)** Since the resulting read datasets represent different enrichments based on the same original community, they are optimal for coassembly as well as co-abundance variation-based binning approaches. An unbiased representation of the source community is achieved by also including the original unsorted sample in the analyses. Created with BioRender.com

## Results and Discussion

### Establishment of midi-metagenomics

In midi-metagenomics, the original sample population is divided into multiple fractions, in which different community members are selectively enriched or depleted **(Figure 2A**). Selective fractionation is achieved *via* fluorescence-activated cell sorting (FACS). Possible strategies for selectively fractionating a complex community into distinct subpopulations are manifold^19^ and can be based on phylogenetic, physiological or morphological properties of the target organisms^20^, e.g., Fluorescent in situ Hybridization (FISH) using rRNA or mRNA probes^11,21,22^, autofluorescence detection^23^ or simply cell size and complexity^24^. In order to improve MAG reconstruction by subsequent co-abundance variation-based binning approaches^25^, fractionation does not need to be particularly stringent as long as a tendential enrichment or depletion can be achieved for at least some of the present taxa. To illustrate this point, community fractionation in this proof-of-principle study was based on relatively simple gating strategies exploiting only those cell characteristics easily detectable *via* FACS: cell size and complexity as defined by forward- and side scatter gating, respectively **(Supplementary Figure S1)**. Since soil represents one the most complex and challenging microbial communities for metagenomic analyses^5,26^, it was chosen as a test environment.

In contrast to standard single-cell and “mini-metagenomics”, approaches which require an amplification step^14,16,20,27,28^, the midi-metagenomic methodology utilizes bulk sorts of several hundred thousand to million cells into the same fraction. However, preliminary trial DNA extractions performed on bulk sorts of bacterial cultures indicated a presence of DNA predominantly in the supernatant and not the pellet^29^ of centrifuged cell suspensions after FACS **(Supplementary Figure S2)**. This observation indicates possible cell damage due to stress, caused by the sorting process, and subsequent release of cellular DNA^29–32^. Therefore, we propose a specially adapted DNA extraction protocol for midi-metagenomic fractions that includes an alcohol precipitation step directly from sorted cell suspensions rather than centrifuged cell pellets, thereby ensuring maximized DNA yields (**Figure 2B**). In preliminary trial runs, DNA yields ranged between 5-30 ng DNA for up to 5 million sorted cells **(Supplementary Table S1)** within sorted midi-metagenomic fractions, which is more than sufficient for direct sequencing^33,34^. Therefore, based on the low input requirements of less than one nanogram^35–37^ for modern sequencing library preparation techniques, sorting efforts may be reduced down to 100 000 cells per sorted fraction for midi-metagenomic sequencing in the future.

In the final practical step of the midi-metagenomic approach, genomic DNA is sequenced seperately for each fraction, as well as the original unfractionated sample **(Figure 2C)**, resulting in multiple individual read datasets. Each of these datasets represents a different composition of the exact same original microbial community, thereby solving a common dilemma for co-assembly as well as co-abundance variation based binning approaches^25,38–40^: Although co-assembly of multiple samples has been shown to increase genome recovery rates especially for low abundant species^41^, it often also produces more fragmented assemblies and increases the risk of strain or species-level chimeras due to increased strain heterogeneity^42^. Such heterogeneities are often introduced by seasonal or locational variability between sample spots or sampling times, which unfortunately is the same variability that is supposed to be exploited by co-abundance variation-based binning approaches.

Since in midi-metagenomics all fractions originate from the same collected sample and thereby same basic community, this approach prevents inter-sample strain variability, hence maintaining optimal conditions for co-assembly as well as binning **(Figure 2D**).

### Efficiency of cell sorting based community fractionation

The relationship between sorted fractions and corresponding unsorted samples was analyzed based on 16S rRNA gene diversity within the assembled metagenomic and midi-metagenomic fractions **(Figure 3, Supplementary Table S2)** as well as 16S rRNA amplicons of increased sequencing depth **(Supplementary Figure S3, Supplementary Table S3**). Weighted UniFrac scores calculated from these analyses show higher beta-diversities between sorted fractions and their respective non-fractionated communities than between non-fractionated samples taken at different years and seasons **(Figure 3)**. This increased beta-diversity represents a strong shift in relative taxon abundances within the respective microbial communities, which can be exploited for distinguishing different organisms based on differential coverage information during downstream binning attempts. At the same time all sorted fractions show decreased alpha-diversity values, and therefore lower community complexity compared to their respective non-fractionated counterparts **(Supplementary Figure S4**).

**Figure 3:**
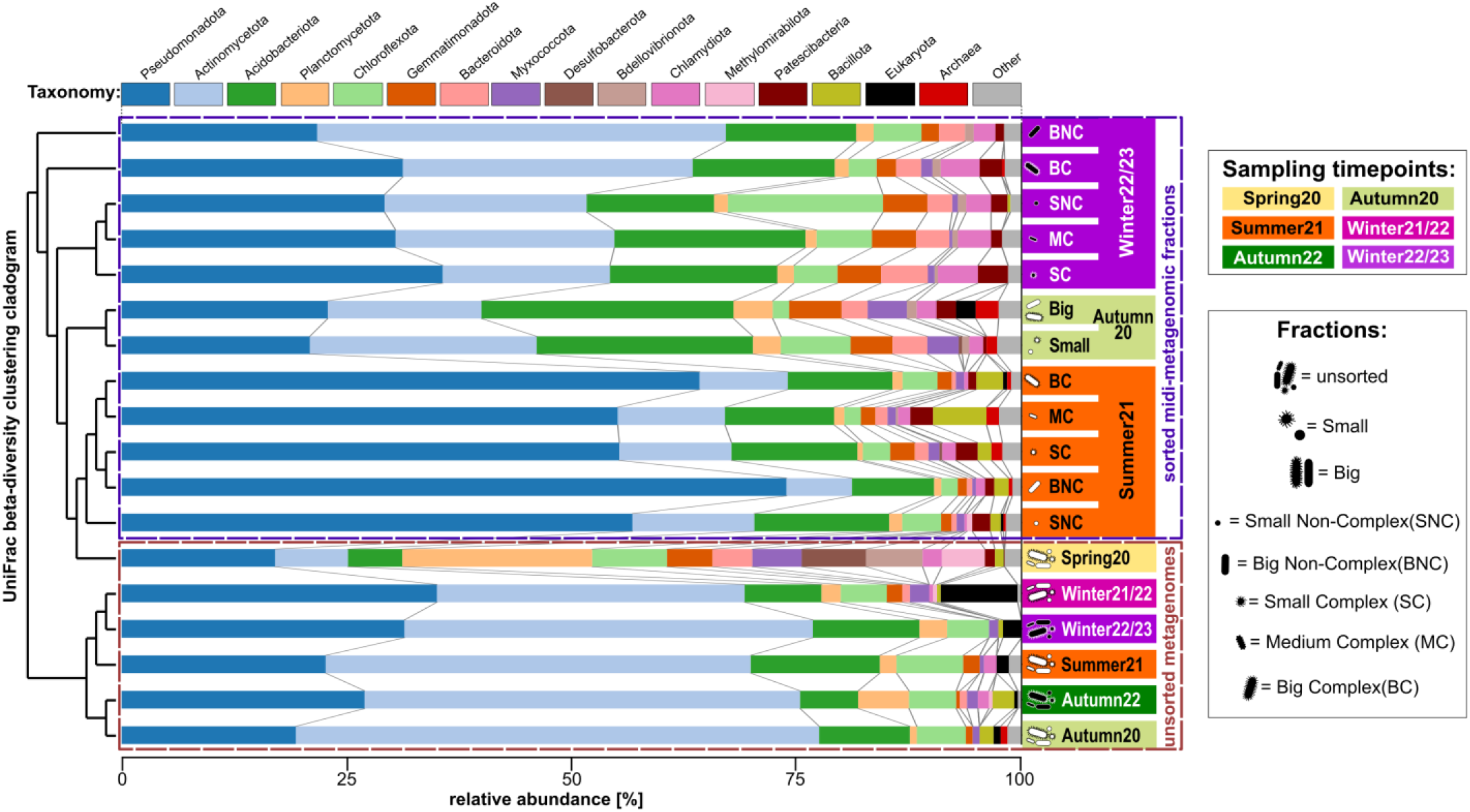
16S rRNA gene based diversity among different (midi-)metagenomic fractions of different samples. Clustering is based on weighted UniFrac^43^ beta-diversity scores and is shown as cladogram on the left. The background colouring of Y-Axis labels on the right side indicates the respective origin-sample. Stacked bar charts indicate the community composition of each sample and fraction, with different phyla being indicated by a distinct colour code as indicated above the plot, and relative abundances being indicated by bar heights according to the X-axis below the plot. Sorted midi-metagenomic fractions are indicated by pictograms and abbreviations as given in the legend on the right.

One sample exclusively sorted on forward scatter signals (which roughly indicate cell size), clusters closer to the unsorted sample when analyzed on high-depth amplicon sequencing level **(Supplementary Figure S3**). This indicates that the use of the forward scatter alone provides a less systematic separation of the community compared to sorting based on forward- and side-scatter combined, possibly due to the reduced resolution of morphological differences. Additional sorting metrics such as fluorescence should therefore even further improve binning efficiency.

### Assembly and binning performance

Furthermore, co-assemblies of standard metagenomics and midi-metagenomics were compared using always the same total sequencing depth of 15 Gbp (averaging at 70 million read pairs per co-assembly) equally distributed across the combined samples and fractions (**Supplementary Table S4**). Based on maximum contig length and N50 metrics, midi-metagenomic co-assemblies of sorted fractions originating from the same original sample were significantly less fragmented than co-assemblies of distinct samples **(Figure 4)**, with p < 0.001 as determined *via* Mann-Withney U tests^44^. This effect is therefore most likely connected to the above mentioned common dilemma in metagenomics: The frequent disparagy between increased binning resolution^41^, but reduced contig sizes when co-assembling multiple datasets, caused by seasonal variability between samples^42^. The improved assembly metrics of the midi-metagenomic approach are therefore likely due to an overall reduced community complexity in the sorted fractions (**Supplementary Figure S4**), but also to the absence of seasonal or regional variability. As a result, the distribution of sequencing efforts over multiple midi-metagenomic fractions maximizes both, coverage variation and sequencing depth in a cost-efficient manner.

**Figure 4:**
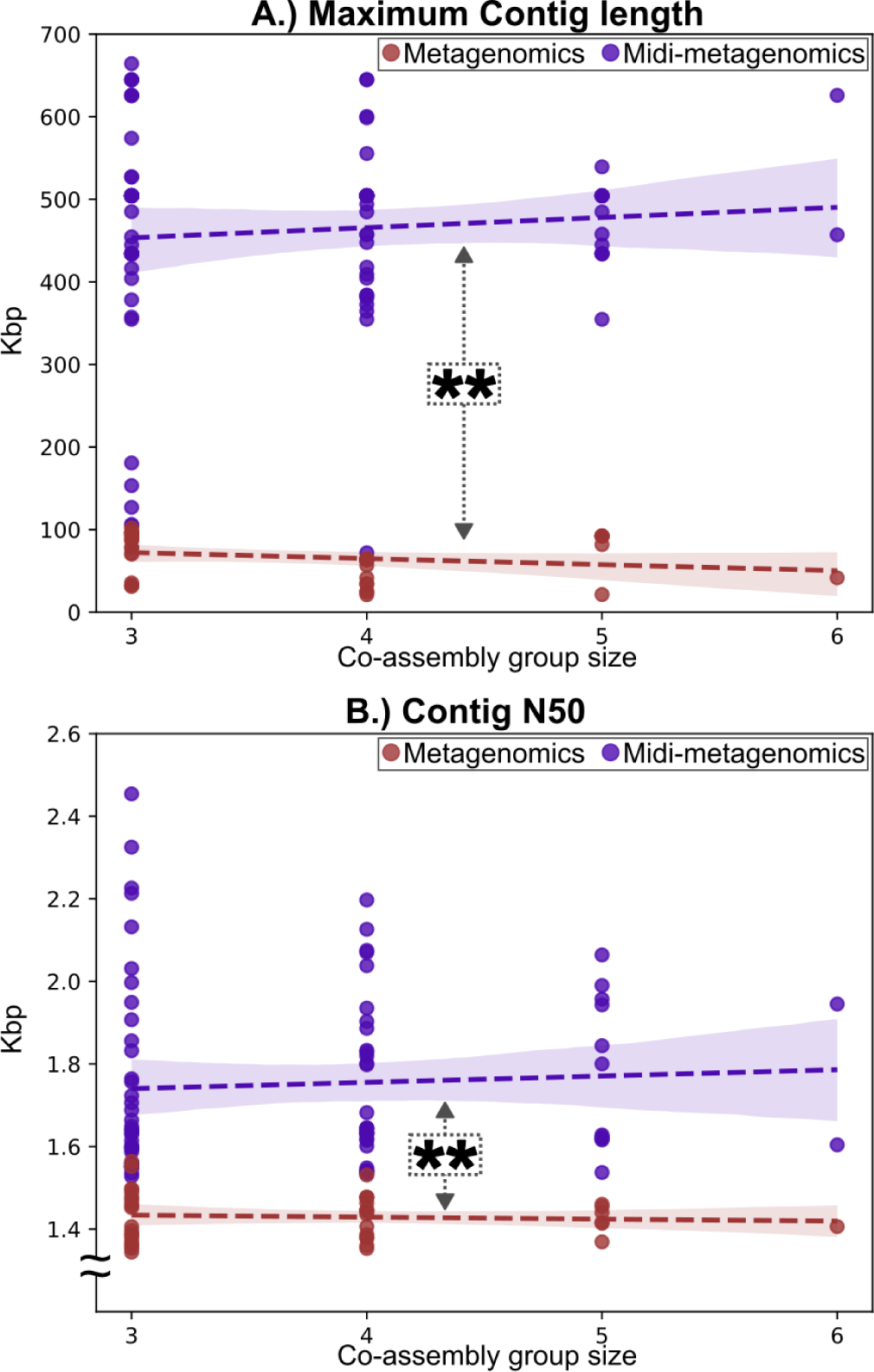
Comparison of assembly metrics for metagenomic and midi-metagenomic approaches in dependance of co-assembled samples and fractions. Scatterplots showing Metagenomic results as red, and Midi-metagenomic results as purple dots. Trendlines and corresponding confidence areas were determined by regression analysis and are indicated by dashed lines and background coloring, respectively. A.) Maximum contig lengths B.) Contig N50 values. The significance of differences in the distribution between Metagenomic and Midi-metagenomic assembly metrics were determined *via* Mann-Whitney U tests^44^

Expectedly, improved assembly metrics also affect the distribution of quality categories among the produced MAGs: Even before contamination filtering with MDMcleaner^45^, midi-metagenomic approaches produced far more high quality MAGs (completeness >90%, contamination <5%) compared to standard metagenomic assemblies, which predominantly consisted of only low quality or moderate quality genomes (completeness >50%, contamination >10%) with contamination values typically above 5% (**Fig 5A-C, Supplementary Table S5**).

**Figure 5:**
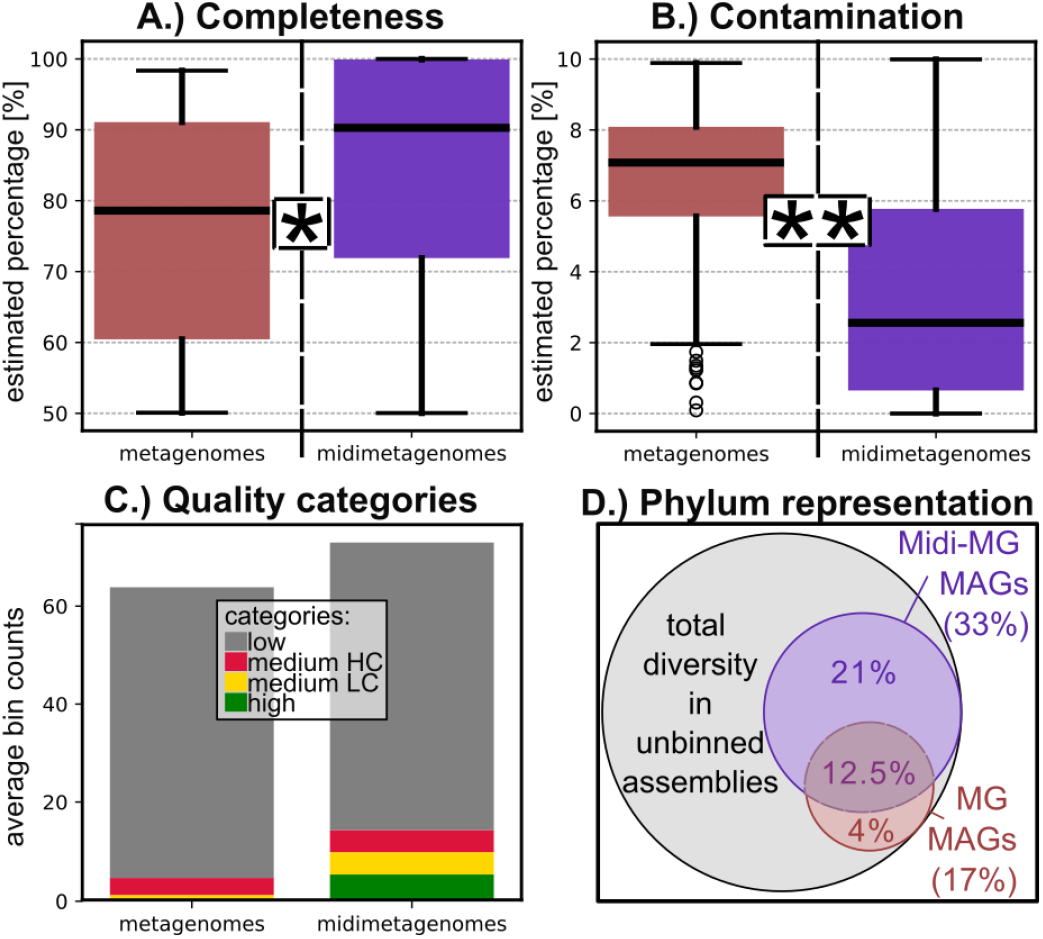
Comparison of quality metrics and diversity of MAGs obtained from standard metagenomic and midi-metagenomic co-assemblies. **A&B.)** Boxplots showing the distribution of checkm2 completeness and contamination estimates, respectively. The difference between metagenomic and midi-metagenomic results was found to be statistically significant (p < .01 based on Moods Median Test^46^) in both cases A more detailed plot showing the individual results for each sample and coassembly group size is given in supplementary Figure S5 **C.)** Average number of MAGs belonging to different quality categories obtained by standard metagenomic and midimetagenomic approaches. **D.)** Relative fractions of total phylum level diversity detected in the unbinned metagenomic co-assemblies that are represented by metagenomic and midi-metagenomic MAGs, respectively.

These trends persist across different midi-metagenomic samples and co-assembly subset groups of different sizes, despite varying community complexities, prooving the robustness of the approach. Interestingly, midi-metagenomic MAGs represented almost twice as many distinct phyla than standard MAGs **(Figure 5D)**, illustrating another important aspect of improved assembly and binning metrics of the midimetagenomic approach: Improved representation of original sample diversity. This is further corroborated by detailed phylogenomic analyses of low contaminated MAGs (<5 % contamination estimate) with at least moderate (50%) completeness (**Figure 6**), which indicate a far broader and, due to increased MAG qualities, also more reliable, phylogenomic representation by midi-metagenomic MAGs compared to standard metagenomics. Interestingly, the midi-metagenomic approach also appears to have reconstructed a higher diversity of closely related but still distinct genomes (as shown in Figure 6 in the case of *Acidobacteriota*, and to a lesser extent also *Alphaproteobacteria* and *Actinomycetota*). This indicates a better resolution of sequence homologies between closely related organisms, by midi-metagenomics compared to standard metagenomics.

**Figure 6:**
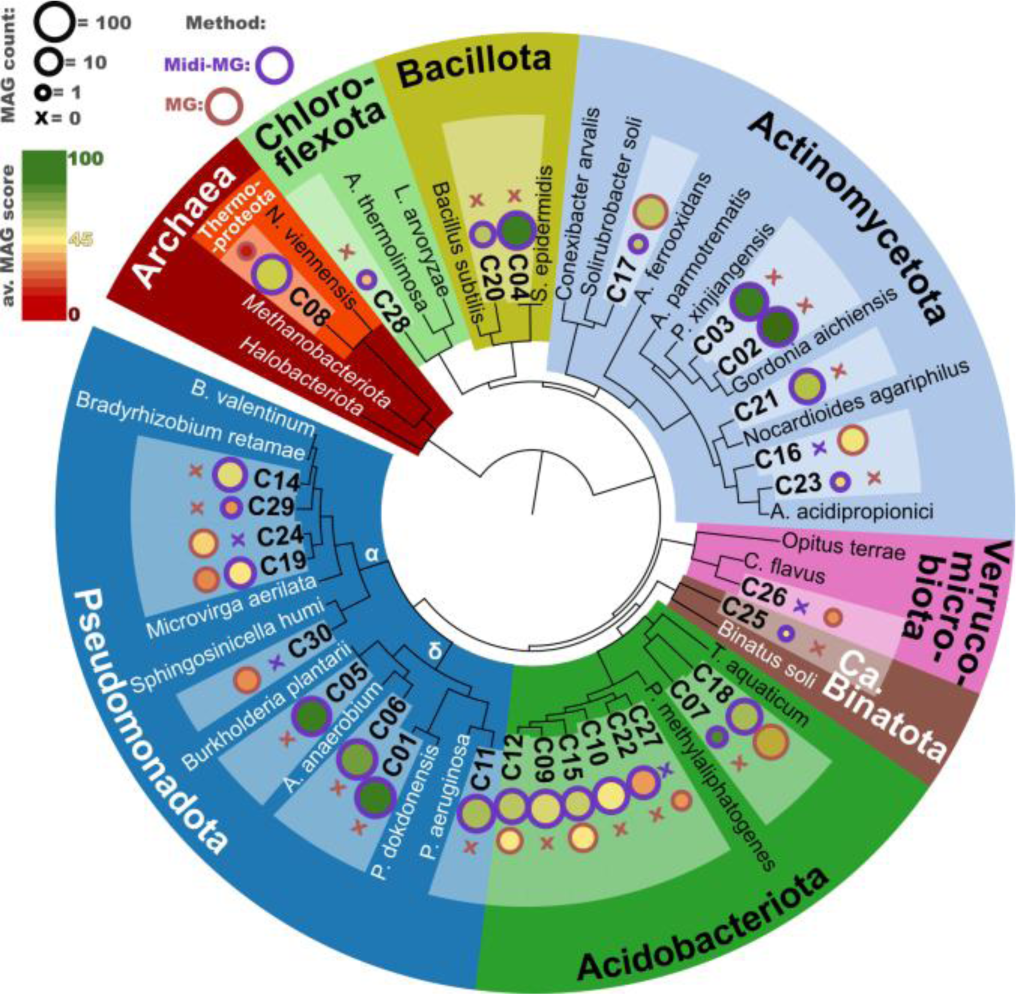
Multi Locus Sequence Analyses (MLSA) based phylogeny of representative MAGs and related reference genomes. Maximum likely-hood phylogenetic clustering based on 61 single copy orthologs shared by all comparison genomes, concatenated to a total length of 7178 amino acids. The software tool dRep^42^ was used to group all MAGs on species level based on ANI comparisons, and to rank the members of each group based on genome quality. Only groups with representatives showing >50% completeness and <=5% contamination were considered, resulting in 30 groups labelled C1-C30, for each of which only the best representative is compared. Bubble Plots next to each group designation indicate the number and average dRep score of each group, as indicated by the legend on the upper left. α = alphaproteobacteria, δ = gammaproteobacterial, MG = Metagenomics, Midi-MG = Midi-meta-genomics

### Comparison with alternative assembly and coverage distribution strategies

While the co-assembly of different fractions theoretically maximizes the potential of the midi-metagenomics approach by optimizing read coverage for assembly and co-abundance based binning, there is some debate within the scientific community over the applicability of this approach for standard metagenomics of multiple distinct samples, due to the afore-mentioned complexities introduced by seasonal or regional variations. In order to present an objective comparison of the performance of midi-metagenomics vs. metagenomics, we therefore tested alternative assembly strategies for the same overall sequencing depth of 15 Gbp (**Supplementary Figure S6**). A common strategy is to perform separate single assemblies for each sample, and then map every read dataset against each individual assembly^42^. This strategy greatly reduced the performance of both, metagenomic and midi-metagenomic approaches, increasing the fraction of “low quality” MAGs and reducing the yield of “moderate” to “high quality” MAGs (**Supplementary Figure S6A**). This is not necessarily surprising, as this strategy severely limits the sequencing depth available for optimal assembly of differentially abundant genomes but also for efficient taxonomic resolution of subsequent binning steps. Interestingly however, although the midi-metagenomic approach completely loses its entire theoretical advantage due to this drawback, it still performed at least as well as standard metagenomics. The general negative effect of the division of available sequence coverage across separate assemblies may be mitigated by increasing overall sequencing efforts. This however, would drastically increase costs and in many cases limit the financial feasibility of incorporating additional samples or fractions.

A cheaper alternative sequence coverage distribution strategy could therefore be to focus sequencing and assembly efforts only on one “main” sample, and supplementing this with additional lower depth “auxiliary” read datasets of other samples or fractions exclusively for mapping and binning purposes. However, when tested on sub-samples of the here analyzed soil datasets, even this could not mitigate the negative effects of the “single assembly” strategy for standard metagenomics, instead only showing a slightly positive effect for midi-metagenomic datasets (**Supplementary Figure S6B**). The co-assembly of midi-metagenomic fractions is therefore not only the most effective, but also more cost-efficient approach since the entire sequencing depth can be fully utilized both for assembly and mapping. A particularly noteworthy observation is the fact that for midi-metagenomics, the co-assembly of “auxiliary” sequencing coverage datasets -despite being less effective than an equal distribution of sequence coverage -still outperformed all standard metagenomics approaches. Thus, the sequencing of an unsorted “main” fraction supplemented by auxiliary sorted fractions is a suitable method for further cost-reduction that maximizes sequencing depth for the original un-fractioned sample community while still enabling optimal binning of the respective MAGs based on less thoroughly sequenced sorted fractions.

### Comparison with mini-metagenomics

We also applied mini-metagenomics to one sample. The mini-metagenomic approach is designed to reduce MDA bias by supplying higher amounts of input DNA. Accordingly, we encountered fewer negative MDA reactions and more complete genomes using this approach compared to standard single cell genomics. Since the advantage of single cell resolution is completely lost with this approach and the binning of the produced contigs is suboptimal without reliable coverage information, only two moderate quality mini-metagenomic MAGs could be recovered, both displaying high contamination estimates close to the MIMAG cut-off of 10% (**Supplementary Table S5, Supplementary Figure S6C**). This indicates a lower cost-efficiency of the mini-metagenomic approach compared to midi-meta-genomics with at least 9x, and potentially even 20x, higher sequencing cost per MAG (**Supplementary Table S6)**.

## Conclusion

Midi-metagenomics produces more diverse and higher quality MAGs compared to classic metagenomics **(Figure 6)** and is also more cost-efficient than mini-metagenomics. While more stringent sorting criteria can be expected to lead to futher improved results, we could show that simple sorting set-ups are sufficient for substantial increase in binning quality as long as partial enrichment or depletion of different community members can be achieved in different fractions. In fact, just the sorting itself already represents a general depletion of large multi-cell aggregates, extracellular DNA as well as potential stress susceptible cell types^29^. Sorting can be based on many different cell properties, of which the here utilized cell size and complexity are only the most simple examples^19^. Possible alternatives could be targeting specific taxonomic groups based on previously established 16S rRNA FISH labelling^21^, or function based enrichments using specific mRNA targeting probes, fluorescent labelled antibodies or even just sorting based on different autofluorescence spectra caused by species specific membrane protein compositions^23^.

An accurate representation of the actual natural community is maintained simply by including an unsorted shotgun metagenome library for each sample **(Figure 2)**. Because this unsorted metagenome can seamlessly be integrated into the analyses as a separate fraction by treating it identically as the sorted fractions during co-assembly and binning steps, the overall effort is not necessarily higher than for standard multi-sample approaches. Consequently, midi-metagenomics may also serve to boost binning efforts in cases where the variability between samples may turn out not be sufficient for co-abundance-based binning, especially for sampling locations that are hard or expensive to access for additional sampling trips, i.e. deep-sea sediments. The exact sorting criteria do not even need to be decided beforehand as a glycerol stock of frozen sample can be revisited for sorting even after preliminary whole-community metagenome analyses.

The most significant advantage of the midi-metagenomics approach, however, is simply the maximization of the available sequence data for assembly as well as co-abundance variation-based binning purposes, while simultaneously avoiding the complications typically introduced by inter-sample strain heterology. This means that read depth may be distributed across multiple fractions without negatively affecting assembly quality (**Figures 4 & 5**), therefore increasing the yield and reducing the required sequencing cost compared to standard metagenomic and mini-metagenomic approaches (**Supplementary Table S6**).

This also ensures a higher sequence and coverage resolution, enabling a more reliable distinguishment between falsely and correctly binned contigs. As a result, there is substantially less likelihood of contamination in the produced MAGs compared to traditional metagenome binning approaches. This is of particular significance as the minimization of MAG contamination desperately needs to be prioritized, considering recent complaints of increasing reference database contaminations caused by insufficiently screened MAGs and SAGs^9,47,48^. Consequently, midi-metagenomics represents a novel improved technique for mining reliable microbial dark matter genomes from environmental samples.

## Acknowledgements

The Authors acknowledge support by the state of Baden-Württemberg through bwHPC. The authors furthermore want to acknowledge the support by Dr. Florian Lenk in proofreading this manuscript and providing helpful suggestions.

## Author contributions

Study conception and design: John Vollmers, Anne-Kristin Kaster; data collection: Maximiano Cassal, John Vollmers, Analysis and interpretation of results: John Vollmers, draft manuscript preparation: John Vollmers, Maximiano Cassal, Funding: Anne-Kristin Kaster

## Materials and Methods

### Microbial Samples

To evaluate midi-metagenomics performance compared to metagenomics, soil samples were collected at the Karlsruhe Institute of Technology (KIT) – Campus North, Eggenstein-Leopoldshafen (49°5’48.8’’N, 8°25’55.6’’E), Germany, during four different periods of time: October 7^th^, 2020, May 25^th^, 2020, August 10^th^, 2021, and February 15^th^, 2022. From each sample, several grams were directly frozen at −80 °C immediately after collection for subsequent standard metagenome DNA extraction and sequencing.

Five grams of each sample was then prepared for cell sorting by adding 30 mL of filtered, autoclaved and UV-sterilized Phosphate Buffer Saline (PBS) solution, brief vortexing to disrupt aggregates and dislocate cells attached to debris, and subsequent pelleting and removal of debris by brief centrifugation at 2,000 × g. Sterile glycerol was added to a final concentration of 30% as an anti-freezing agent and the samples were stored at −80°C until further processing. An overview of all samples is given in **Table 1**.

**Table 1:** Overview of samples and fractions. Fraction abbreviations: BC = “Big Complex”, MC = “Medium Complex”, SC = “Small Complex”, BNC = “Big Non-Complex”, SNC = “Small Non-Complex”,

| Sample | Sampling location | Sampling date | Fractions produced |
| --- | --- | --- | --- |
| spring20 | KIT, Campus North<br>49°5′48.8″N,<br>8°25′55.6″E | 15.05.2020 | only **unsorted** |
| autumn20 |  | 07.10.2020 | **unsorted, big & small** |
| summer21 |  | 10.08.2021 | **unsorted, BC, MC, SC, BNC & SNC** |
| Winter21/22 |  | 15.02.2022 | **unsorted, BC, MC, SC, BNC & SNC** |
| Autumn22 |  | 07.10.2022 | only **unsorted** |
| Winter22/23 |  |  | **unsorted, BC, MC, SC, BNC & SNC** |

### Fluorescence-Activated Cell Sorting (FACS)

Prior to sorting, the samples aliquoted for midi-metagenomics were centrifuged for 1 min at 15,871 × g and 20 °C. The supernatant was discarded and after resuspension of the pellet in 1 mL PBS, 5 µl SYBR^®^ Green I was added to all samples. The samples were then vortexed, incubated for 20 min at 4 °C and subsequently pelleted again by centrifugation for 1 min at 15,871 × g. Each pellet was then washed twice with 1 mL PBS.

Before loading the sample into the FACS (BD FACSMelody™, Becton, Dickinson and Company, New Jersey, USA), an unlabeled negative control was filtered into a 5 mL FACS tube using a sterile SYSMEX CellTrics^®^ filter with 20 µM mesh size and then diluted with PBS. The negative control was used to compare the difference of fluorescence signals for a correct gating that included only labelled cells. Subsequently, the same procedure was applied to the SYBR-labelled samples. A threshold was set up in order to disregard smaller particles such as debris during the sorting process and an excitation wavelength of 488 nM was used.

For samples “summer 21”, “winter21/22” and “winter22/23” cells were sorted into five different groups via gatings based on plotting fluorescence intensity against the Forward Scatter Signal (FSC) and Side Scatter Signal (SSC), which are roughly proportional to cell size and complexity, respectively. (**Supplementary Table S1 & Supplementary Figure S1**). For sample “autumn20” only two groups were sorted, according to size measured by differences in FSC (**Supplementary Table S1**). After sorting, the cells were stored at −80 °C until further processing. An overview of the Fractions produced per sample is included in **Table 1**.

### DNA Extraction

For metagenomics of the unsorted sample, DNA was extracted with the Dneasy PowerSoil Kit (Qiagen, Hilden, Germany) following the manufacturer’s instructions. For midi-metagenomics community fractions, DNA was extracted directly from FACS sorted cell suspensions consisting of 4 × 10^6^ cells. First, the cells were freeze-thawed three times using liquid nitrogen and a 60 °C water bath. Then, bead beating was performed three times for 30 s at 6 m/s using one tube of lysing matrix for each fraction (Cat.#6914-800, MP Biomedicals, Ohio, USA) and an MP Bio Fast Prep^®^-24 homogenizer (MP Biomedicals, Ohio, USA). Beads and cell debris were pelleted by centrifugation at 14,000 ×g for 5 min and the supernatant was subjected to standard alcohol precipitation using 1 volume of 80% isopropanol, 0.1 volume 3 M Sodium Acetate and 340 µg Linear Polyacrylamide. After a subsequent wash step with ice cold 70% ethanol the resulting DNA pellet was resuspended with 100 µl PCR-grade water followed by further purification *via* solid-phase reversible immobilization using 1.5 volume of AMPure XP Beads (Beckman Coultier™) and final elution in 20 µl 1× TE. All extracted DNA was immediately stored at −20 °C until use.

### Polymerase Chain Reaction (PCR) for Amplicons

Amplicon sequencing was performed using a nested PCR approach. Almost full-length PCR products were obtained in a preliminary PCR using 1.25U OneTaq^®^ Quick-Load^®^ DNA Polymerase (New England BioLabs, Ipswich, MA, USA), 200 µM mixed dNTPs, 500 µM biology-grade Bovine Serum Albumin (BSA) (Thermo Fisher Scientific, Waltham, Massachusetts, USA) and 0.2 µM of each universal bacterial forward and reverse primer 27F (5’-AGRGTTYGATYMTGGCTCAG-3’) and 1492R(5’-AGRGTTYGATYMTGGCTCAG-3’). PCR products were purified using DNA Clean & Concentrator™-5 columns (Zymo Research Europe GmbH, Irvine, California, USA) according to the manufacturer’s instructions. The purified product was then used as template for a subsequent amplicon PCRs using 0.5 U Q5^®^ High-Fidelity DNA Polymerase (New England Biolabs, Ipswich, MA, USA) 0.5 U, 200 µM dNTP Solution Mix (New England Biolabs), Q5^®^ High GC Enhancer, 0.1 µg/µl BSA (Thermo Fisher Scientific, Waltham, Massachusetts, USA) and 0.2 µM of each universal bacterial primer 341F (5’-AGRGTTYGATYMTGGCTCAG-3’) and 518R (5’-AGRGTTYGATYMTGGCTCAG-3’), targeting the V3 hypervariable region.

### Sequencing

All libraries were prepared using the NEBNext® Ultra™ II FS DNA Library Prep Kit for Illumina^®^ (New England Biolabs, Ipswich, MA, USA), according to the manufacturer’s instructions. Libraries were sequenced on an Illumina NextSeq 550^®^ (New England Biolabs, Ipswich, MA, USA) device using 300 cycles and a paired-end approach.

### Read processing and assembly

Reads were quality trimmed and adapter-clipped using trimmomatic v.0.36, bbduk v.35.69 and cutadapt v.1.14 successively^49–51^:. Overlapping read pairs were identified and merged using FLASH v.1.2.11^52^. For amplicon datasets, reads were clustered into Amplicon Sequence Variants (ASV) using the Qiime2 pipeline with dada2. To account for different sequencing depths the produced AVS were rarefied to a value of 4000 which was determined via preliminary rarefaction analysis. Rarefied ASVs were subsequently taxonomically classified using SINA v1.7.2^53^. Shotgun datasets were arranged into co-assembly groups representing all possible subsets of three to six datasets per metagenome or midi-metagenome (**Supplementary Table S4**). In order to enabble standardized assemblies with 15 Gbp total read input each, shotgun datasets were randomly subsampled down to 2.5, 3.75, 3 and 5 Gbp when possible, and coassembled at equal amounts for each co-assembly group. Additional assemblies with non-uniform sequencing depth distribution were also performed using unsorted metagenome samples as “main” datasets with 12-13 Gbp sequencing depth, and 5 “auxilliary” datasets subsampled to 0.4-0.5 Gbp each. Assemblies were performed using MegaHit v1.2.9 ^54^.

### MAG reconstruction and analyses

For each co-assembly, three different binning tools were used in parallel: Metabat2 v.2.15, Concoct and Rosella v.0.4.1^18,39,55^. For Midi-metagenomic approaches, Rosella was substituted for Maxbin^56^, as Rosella did not function without co-abundance information and Maxbin utilizes additional taxonomic and marke-gene criteria that may optimize results for mini-metagenomic and single cell genomic assemblies that lack reliable coverage information ^3,17^. Resulting bins were pre-assessed and filtered using MDMcleaner. Quality categories were then determined based on re-assessments using checkm2^57^. Taxonomic classifications were based on GTDB-TK v2.1.1^58^

dRep v.3.4.0 ^42^ was employed to identify groups of redundant MAGs created by different assemblies or binning tools and to select the respective most representative MAG. Similarities between MAGs were additionally determined and visualized based on gene-content as previously described elsewhere^59^.

## Data availability

All raw sequencing data are available at NCBI under bioproject PRJNA900514. However, to minimize data redundancy and to avoid database corruption, only dereplicated high quality MAGs (highest quality representative of each dRep-cluster depicted in **Figure 6**) and associated coassemblies were uploaded to NCBI. The complete set of assemblies and MAGs, including low quality MAGs with completeness estimates of at least 50% and contamination estimates below 25% can be accessed via zenodo under the DOI: 10.5281/zenodo.13150466

