## Supplementary Figure for "“Midi-metagenomics”: A novel approach for cultivation independent microbial genome reconstruction from environmental samples"

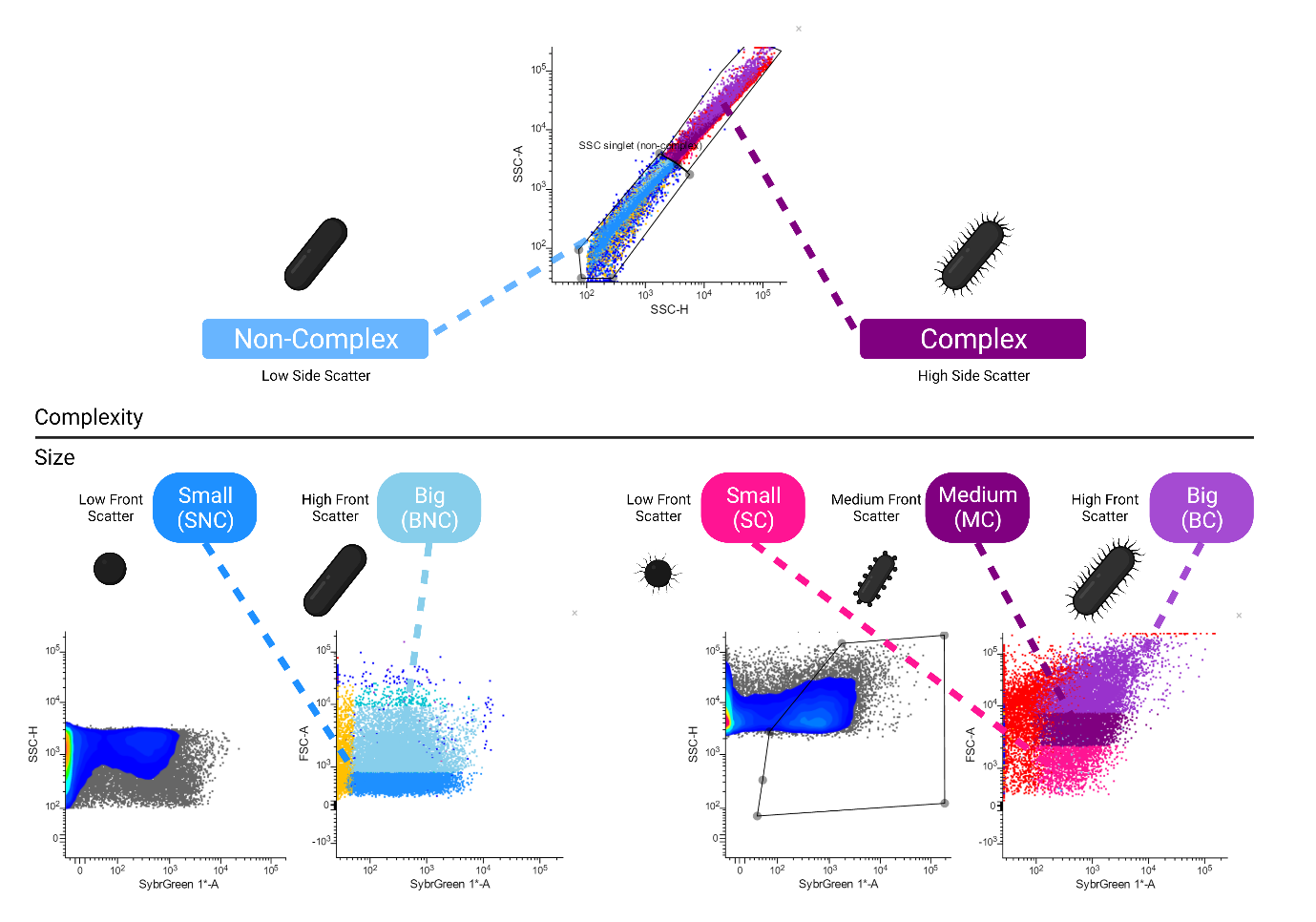

**Supplementary Figure S1: Fluorescence-activated cell sorting (FACS) separation of soil sample into five fractions, according to their complexity and size, discriminated by Side Scatter and Forward Scatter Signals, respectively**.

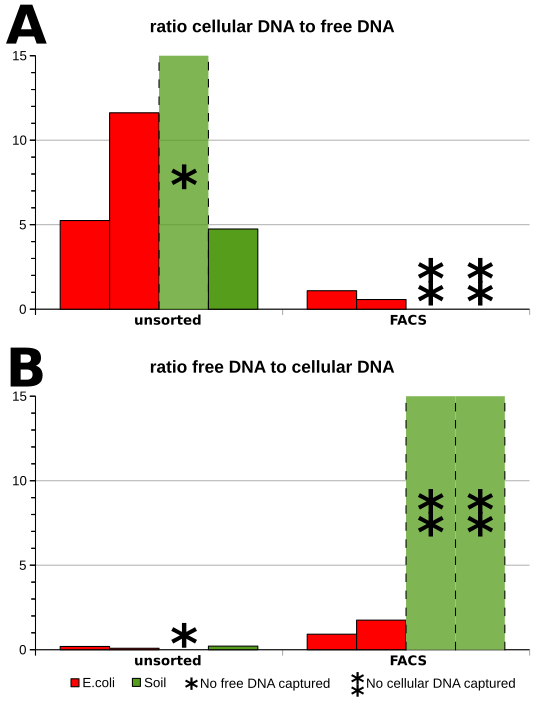

**Supplementary Figure S2**. **Ratio of DNA obtained from cell pellets to DNA obtained from supernatant and vice versa, before and after bulk sorting via FACs**

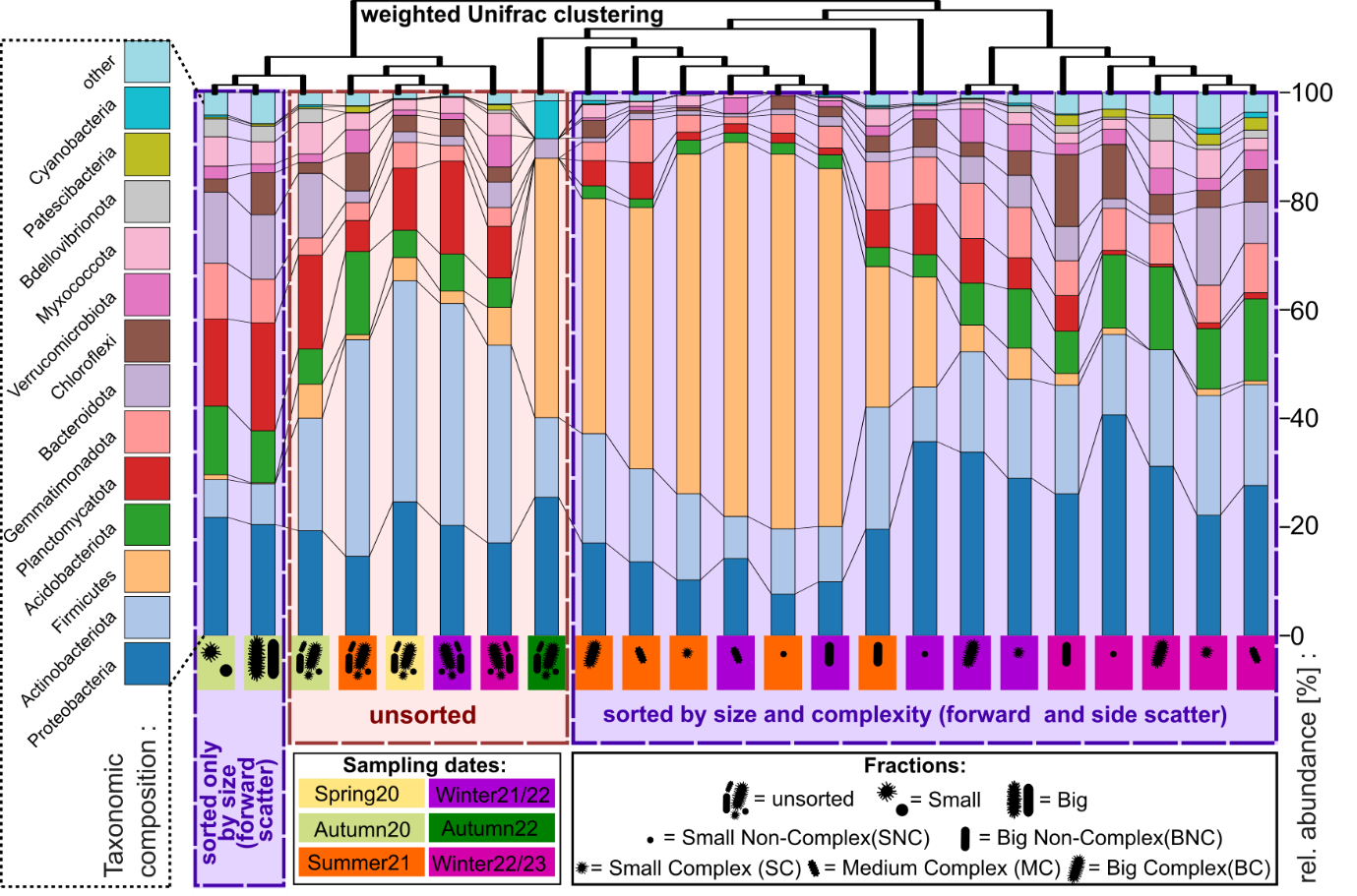

**Supplementary Figure S3:** **Differences in diversity between distinct unsorted samples as well as corresponding sorted fractions, determined based on amplicons of the 16S rRNA V3-region.** Clustering is based on weighted unifrac betadiversity scores and is shown as cladogram. Samples are indicated by background colouring, while fractions are indicated by pictograms, according to the legend at the bottom. Stacked bar charts indicate the community composition of each sample and fraction, with different phyla being indicated by a distinct colour code as indicated on the left. For the “Spring20”and “Autumn22” sample, only the unsorted complete community was analyzed.

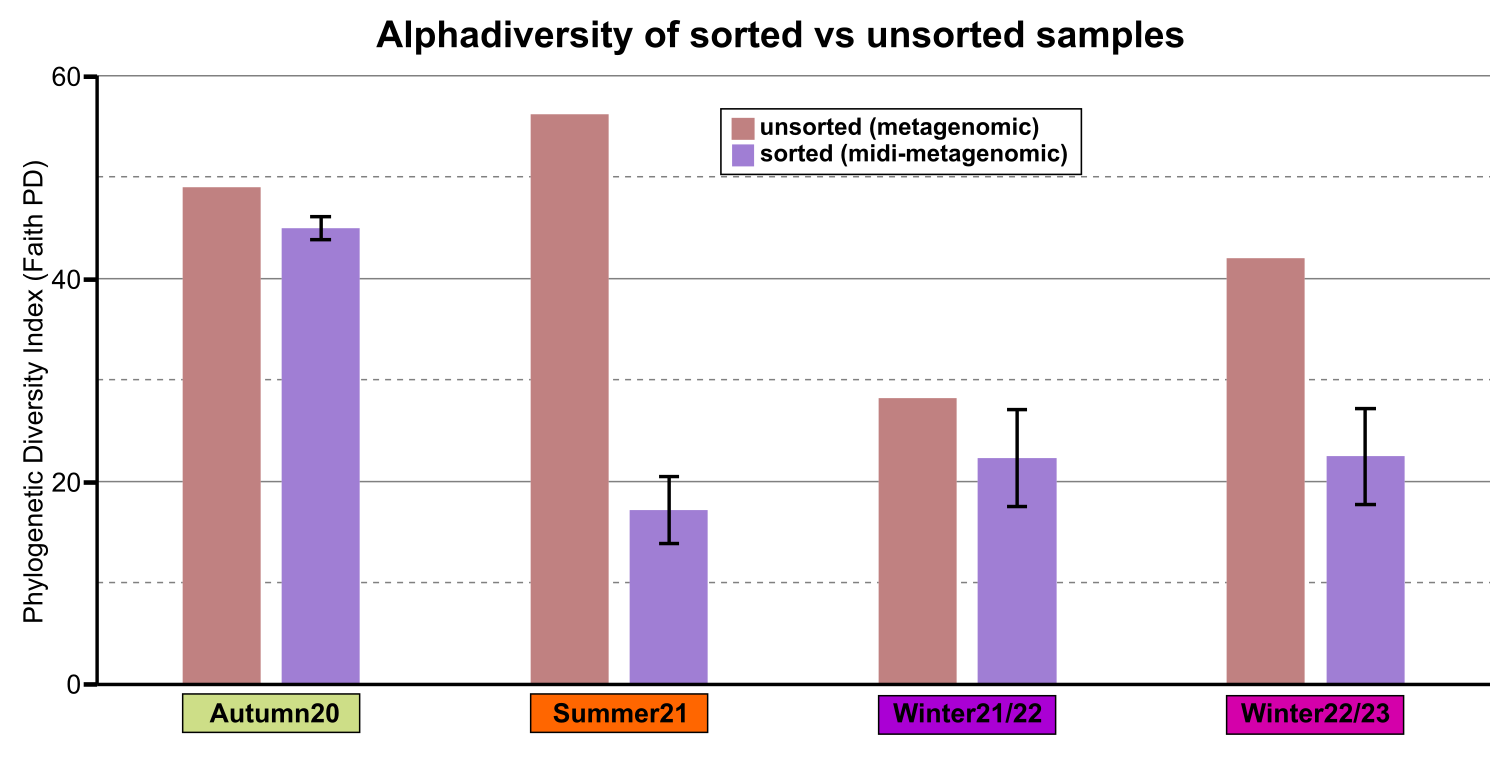

**Supplementary Figure S4: Alpha-diversity comparison between 16S rRNA gene amplicons of different metagenomic samples and corresponding sorted fractions**. Alpha diversity was determined as Faith PD index. For each Soil sample (Autumn20, Summer21, Winter21/22, Winter22/23), the alphadiversity of the unsorted sample (light red) is plotted against the corresponding sorted metagenome (light purple).

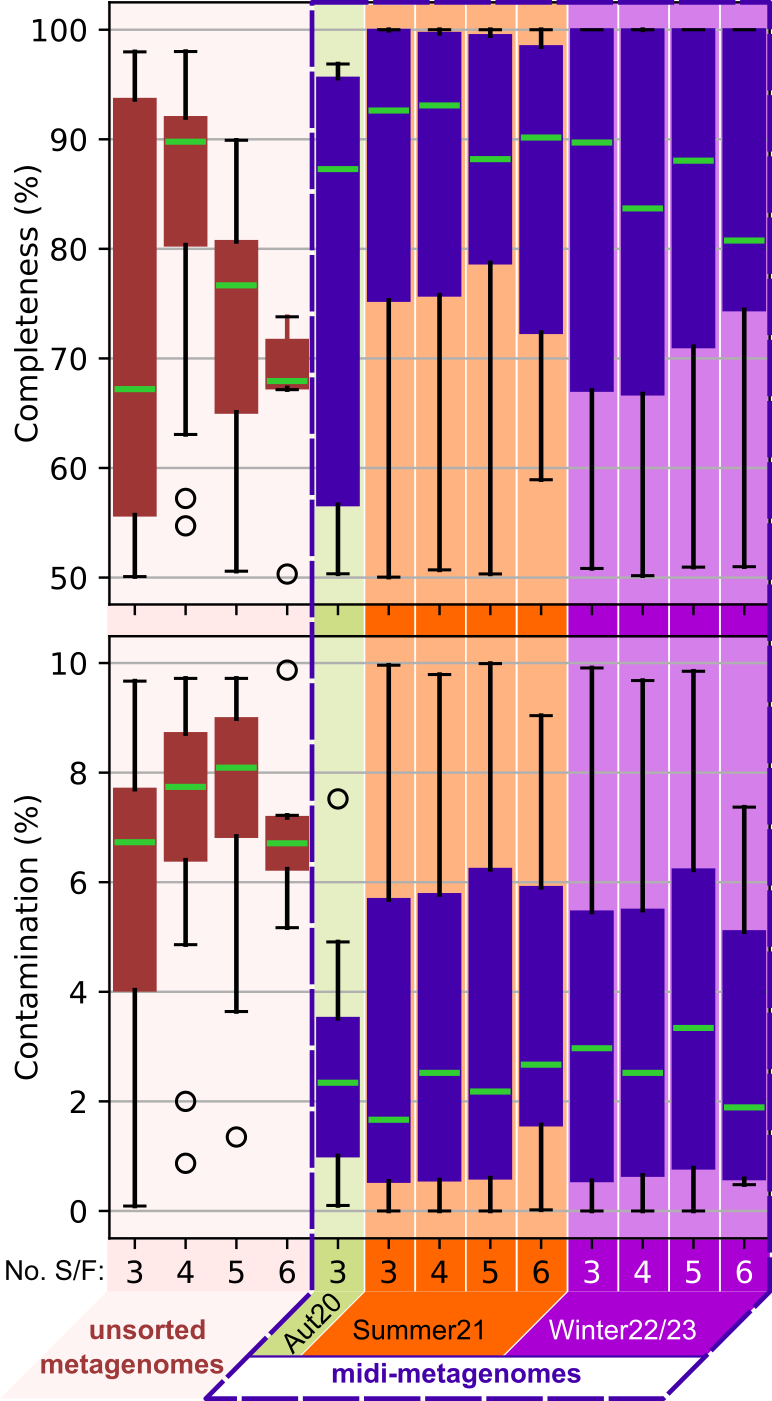

**Supplementary Figure S5: Checkm2 quality metrics of MAGs obtained from standard metagenomic and midi-metagenomic co-assemblies, separated by sample for midimetagenomic Approaches.** Low quality MAGs, as defined according to MIMAG standards (less than 50% completeness or more than 10% contamination) were discarded before analyses. The upper boxplots show the distribution of checkm2 completeness estimates , while the lower plots show the distribution of contamination estimates of all MAGS of moderate quality or better according to MIMAG standards. The number of distinct samples or fractions involved in the respective co-assemblies is indicated on the x-axis (“No. S/F”). Plots for metagenomes are indicated by dark red fill color, plots for midi.metagenomes by dark blue fill color and surrounding box. The data for midi-metagenomes is shown separated by sample, with different background colors indicating different samples. Aut20 = Autumn20. Differences between midi-metagenomic and metagenomic approaches are statistically significant based on Moods median test with p < 0.01.

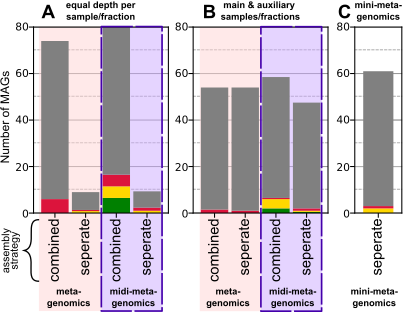

**Supplementary Figure S6: Effect of different strategies for general approach, assembly and sequence depth distribution on binning results.** Subfigures **A & B** show the results for different sequencing depth distribution strategies on the standard- and midi-metagenomic approaches:”equal” and ”main & auxilliary”, respectively. In the ”equal” strategy, sequencing depth was distributed equally over the individual samples and/or fractions. In the ”main & auxilliary” strategy, one (unsorted metagenomic) sample was selected as ”main” sample with ~13 Gbp sequencing depth while the other samples/fractions were threated as ”auxiliary” datasets with only 0.4-0.5 Gbp sequencing depth. Two alternative assembly strategies are also compared: ”combined” were all datasets are combined and co-assembled, and ”seperate” where each (”main”) dataset is assembled individually and all other (”auxiliary”) datasets are only used for mapping. Subfigure **C** shows the result of the”mini-metagenomics” approach.

Colors indicate the MIMAG based quality category of the MAGs: grey = low quality; red = moderate quality (high contamination); yellow = moderate quality (low contamination; green = high quality
